# MORFEE: a new tool for detecting and annotating single nucleotide variants creating premature ATG codons from VCF files

**DOI:** 10.1101/2020.03.29.012054

**Authors:** Dylan Aïssi, Omar Soukarieh, Carole Proust, Beatrice Jaspard-Vinassa, Pierre Fautrad, Manal Ibrahim-Kosta, Felipe Leal-Valentim, Maguelonne Roux, Delphine Bacq-Daian, Robert Olaso, Jean-François Deleuze, Pierre-Emmanuel Morange, David-Alexandre Trégouët, on behalf of the GENMED Consortium

**Author notes:** These authors equally contributed to this work (Dylan Aïssi, Omar Soukarieh).

## Abstract

**Summary:** Variants in 5’UTR regions that create upstream translation initiation AUG codons are a class of neglected non coding variations. When they associate with a premature stop codon and create upstream open reading frames (uORFs) whose translation competes with that of natural proteins, they can have strong impact on human diseases. We here describe MORFEE, a new bioinformatics tool that detects, annotates and predicts, from a standard VCF file, the creation of uORF by any 5’UTR variants on uORF creation. MORFEE was applied to two genomic resources and identified candidate functional variants that could explain statistical association signals observed in the context of Genome Wide Association Studies or could be responsible for rare forms of diseases. In conclusion MORFEE is an easy-to-use tool complementary to existing ones that can help resolving genetic investigations that remained so far unfruitful.

**Availability and implementation:** MORFEE is written in R with code and package available at https://github.com/daissi/MORFEE.

## 1 Introduction

This work was motivated by our recent discovery of a never reported (c.-39C>T) single nucleotide variant (SNV) in the 5’UTR region of *the PROS1* gene that creates an upstream translation initiation AUG codon (uAUG) (Labrouche-Colomer *et al*., submitted). The latter generates an overlapping open reading frame that inhibits the production of the natural Protein S and causes familial inherited Protein S deficiency (Labrouche-Colomer *et al*., submitted). As this variant is located in the *PROS1* promoter region, it was not predicted to be deleterious by popular bioinformatics prediction tools, including PolyPhen (http://genetics.bwh.harvard.edu/pph2/), SIFT (http://sift.bii.a-star.edu.sg/), CADD (Rentzsch *et al*., 2019), REVEL(Ioannidis *et al*., 2016), TraP (Gelfman *et al*., 2017) or RegulomeDB (Boyle *et al*., 2012), most of them mainly dedicated to coding SNVs. Even the PreTIS software (Reuter *et al*., 2016) dedicated to 5’UTR variants only predicted a “moderate” impact of the mutation. We then realize that there was no easy-to-use tool that can automatically read VCF files and detect/annotate SNVs that potentially create premature ATG codon and upstream open reading frame (uORF).

At least 2 situations can result from the creation of a uAUG by a SNV (uaugSNV) located in a 5’UTR region. In the first case, the created uAUG could be in frame with the main coding sequence (CDS) with no associated premature stop codon, likely producing a longer protein. In the second case, the 5’UTR variant could create a uORFUpstream open reading frames are sequences formed by a uAUG, associated with a stop codon preceding the end of the main CDS. The stop codon could be located in the 5’UTR or within the main CDS, resulting in a fully upstream or an overlapping uORF, respectively. Even if the translational role of uORFs still remains unclear, it is known that these elements reduce the production of the main protein (Barbosa *et al*., 2013). Furthermore, uORF creating variants (referred to in the following as *loorf* SNV because their potential impact mimic that of loss-of-function (lof) coding variant) could alter the protein expression and could be disease-causing variants (Romanelli Tavares *et al*., 2019). The effect of uORF on the main protein expression depends on many elements, including the uORF length (longer uORFs are hypothesized to associate with a higher level of downstream translation repression), the uORF-CDS distance (shorter distance between the upstream stop codon and the CDS start could be associated with lower levels of the main protein) and the kozak context (Wethmar, 2014). Information about these elements are of key importance to predict as accurate as possible the impact of a *loorf* at the protein level. In absence of an automated method, this type of predictions is very time-consuming and not adapted to the huge number of variants identified by new generation sequencing nor to the large variety of uORF.

To fill this gap, we develop a R program, MORFEE (for Mutated upstream ORF dEtEction), that detects, in a given VCF file, any 5’UTR SNV that could potentially create premature ATG codon and provides some annotation about their potential impact on the natural protein. The application of MORFEE to two genetic datasets illustrates that *loorf* are a neglected class of non-coding variations that could contribute to human diseases.

## 2 Implementation and application

### 2.1 Workflow

The algorithm of MORFEE is written in R language and can run on all operating systems which have an R interpreter (including Linux, macOS and Windows).

MORFEE starts with a minimal VCF file (i.e. with at least the *chr*, *position*, *reference allele* and *alternate allele* fields) as an input, but can also work with already ANNOVAR-annotated (Wang *et al*., 2010) VCF files. MORFEE has some R packages dependencies which are available on CRAN or Bioconductor repositories and uses the GENCODE database.

In a first step, MORFEE reads the input VCF file and use ANNOVAR (that has then to be beforehand installed) through the wrapper function *vcf.annovar.annotation* to annotate all variants. This step is skipped if the input file has already been annotated. Then, 5’UTR variants associated to any transcript are selected and GENCODE database is used to extract necessary transcripts information to generate the complementary DNA sequence at selected variant. From this sequence, MORFEE determines whether the variant creates a new upstream ATG sequence. When a new start codon is created by a variant, MORFEE calculates: 1-the distance between the reference ATG and the new ATG; 2-the reading frame shift (0 for in frame variant, 1 or 2 for a shift of one or two nucleotides); and 3- the predicted length of the uORF (i.e. the distance between the new ATG and the first stop codon in frame). Finally, a new annotation MORFEE field is added to VCF file with the information about the new ATG. To facilitate the reading of MORFEE results, a second outputfile is generated into a Open XML file (.xlsx) format and including only the potential detected uaugSNVs.

### 2.2 Results

We first applied MORFEE to the 1000Genomes reference dataset (phase 3 v.20130502) and identified 5,161 uaugSNVs among which 4,639 could be *loorfs* (Supplementary Table S1). Among these candidate *loorfs,* 107 are mapping to loci associated with clinical disorders as reported in ClinVar (version 20190305) or to loci identified in Genome Wide Association studies (GWAS). These observations warrant further deep investigation to determine whether such *loorfs* could contribute to explain GWAS signal. For instance, the *FRMD5* rs492571 is in strong linkage disequilibrium with several SNPs reported to associate with triglycerides and HDL-cholesterol levels (Teslovich *et al*., 2010), all these SNPs being considered as intronic. Interestingly, 7 protein coding transcripts, harbouring different CDS and 5’UTR, are associated with the *FRMD5* gene. The rs492571 SNP is located into introns in 6 over these isoforms, but in the 5’UTR of the transcript NM_001286491.2, at c.-487 position. Based on MORFEE predictions, the A>G substitution at this position (rs492571), creates a new start codon in the 5’UTR of the NM_001286491.2 isoform, thus generating a fully upstream uORF carrying 35 codons (including the stop codon; i.e. 34 amino acid). These data suggest that the rs492571 could be a good culprit candidate for the observed GWAS signal whose functionality deserve to be further investigated.

MORFEE was also applied to the vcf file derived from the whole genome sequencing analysis of 200 unrelated patients with idiopathic venous thrombosis (VT) from the MARTHA study (Oudot-Mellakh *et al*., 2012) (Supplementary Materials). Full results listing all identified uaugSNV are given in Supplementary Table S2. MORFEE identified 1,096 uaugSNVs containing 702 uncommon variants (minor allele frequency < 1% or never been reported). Among them, 635 were loorfs, including two variants mapping to candidate VT genes: *PRO*C rs116169054 and *PLAT* rs763007802. MORFEE predictions indicate that the *PROC* SNP rs116169054 could generate an overlapping 35 codons uORF by creating a uAUG in the 5’UTR of the transcript NM_000312.4 while the *PLAT* rs763007802 is predicted to create a uAUG in the 5’UTR of the transcript NM_000930.5, resulting in the production of a 3 codons uORF. Preliminary experimental work (Supplementary Materials) suggest that the presence of the *PROC* variant, but not that of the *PLAT* variant, is associated with decreased levels of the corresponding protein (Supplementary Fig 1.). The effect of the *PROC* rs116169054 *loorf* on decreased protein C levels could be compatible with this variant contributing to VT risk, protein C deficiency being a major risk factor for VT. However, it is not our intention to claim that this *loorf* causes VT but rather to illustrate a case where MORFEE identifies *loorfs* with biological effects compatible with the studied disease. Conversely, the *PLAT* rs763007802 is unlikely to be causal as it does not seem to impact on protein levels. This observation is consistent with what is generally admitted for fully upstream ORF of small length that are expected to have little biological impact. Indeed, the translation reinitiation capacity by the ribosomes at the main AUG is easier with fully upstream uORFs compared to overlapping uORFs (Wethmar, 2014). Moreover, the generated PLAT uORF length corresponds to the shortest potential uORF of 9 nucleotides (Calvo *et al*., 2009), generally associated with a low probability of protein repression.

## 3 Conclusion

MORFEE is an easy-to-use R program that ables, from standard VCF files, to detect and annote SNVs potentially creating premature ATG codon. Its application to two sequence datasets illustrates the relevance to further investigate such neglected type of SNVs, both in the context of common and rare forms of human diseases.

## References

Barbosa,C. et al. (2013) Gene expression regulation by upstream open reading frames and human disease. PLoS Genet., 9, e1003529.

Boyle,A.P. et al. (2012) Annotation of functional variation in personal genomes using RegulomeDB. Genome Res., 22, 1790–1797.

Calvo,S.E. et al. (2009) Upstream open reading frames cause widespread reduction of protein expression and are polymorphic among humans. Proc. Natl. Acad. Sci. U.S.A., 106, 7507–7512.

Gelfman,S. et al. (2017) Annotating pathogenic non-coding variants in genic regions. Nat Commun, 8, 236.

Ioannidis,N.M. et al. (2016) REVEL: An Ensemble Method for Predicting the Pathogenicity of Rare Missense Variants. Am. J. Hum. Genet., 99, 877–885.

Labrouche-Colomer,S. et al. (submitted) A novel rare c. −39C>T mutation in the PROS1 5’UTR causing PS deficiency by creating a new upstream translation initiation codon and inhibiting the production of the natural protein. BIORXIV/2020/007328.

Oudot-Mellakh,T. et al. (2012) Genome wide association study for plasma levels of natural anticoagulant inhibitors and protein C anticoagulant pathway: the MARTHA project. Br. J. Haematol., 157, 230–239.

Rentzsch,P. et al. (2019) CADD: predicting the deleteriousness of variants throughout the human genome. Nucleic Acids Res., 47, D886–D894.

Reuter,K. et al. (2016) PreTIS: A Tool to Predict Non-canonical 5’ UTR Translational Initiation Sites in Human and Mouse. PLoS Comput. Biol., 12, e1005170.

Romanelli Tavares,V.L. et al. (2019) Craniofrontonasal Syndrome Caused by Introduction of a Novel uATG in the 5’UTR of EFNB1. Mol Syndromol, 10, 40–47.

Teslovich,T.M. et al. (2010) Biological, clinical and population relevance of 95 loci for blood lipids. Nature, 466, 707–713.

Wang,K. et al. (2010) ANNOVAR: functional annotation of genetic variants from high-throughput sequencing data. Nucleic Acids Res., 38, e164.

Wethmar,K. (2014) The regulatory potential of upstream open reading frames in eukaryotic gene expression. Wiley Interdiscip Rev RNA, 5, 765–778.

