## Supplementary Methods for "MORFEE: a new tool for detecting and annotating single nucleotide variants creating premature ATG codons from VCF files"

Supplementary Materials

*MARTHA patients selection*

The MARTHA study is a population of venous thrombosis (VT) patients recruited at the La Timone Hospital, Marseille (France). Detailed description of the MARTHA study can be found in (Germain *et al.*, 2011; Oudot-Mellakh *et al.*, 2012). All patients were unrelated Caucasian patients with a personal documented history of VT (deep vein thrombosis and/or pulmonary embolism) in the absence of a major VT risk factor (natural coagulation inhibitor deficiency, homozygosity for factor V Leiden or for factor II G20210A mutation, antiphospholipid syndrome) . All VT events were documented by venography, Doppler ultrasound, spiral computed tomographic scanning angiography, and/or ventilation/perfusion lung scan. The date of occurrence of every episode of VT and the presence of precipitating factors (such as surgery, trauma, prolonged immobilization, pregnancy or puerperium, and oral contraceptive intake) were collected. VT was classified as provoked when occurring within 3 months after exposure to exogenous risk factors, including surgery, immobilization for 7 days or more, oral contraceptive use, pregnancy and puerperium, trauma of the lower limb, long travel (by car > 10 hours, by plane > 5 hours. In the absence of these risk factors, VT was defined as unprovoked.

From this sample of VT patients, 200 were selected for whole genome sequencing. These patients were selected to have experienced unprovoked VT. Besides, these patients should have family history of VT or multiple unprovoked VT events, such clinical patterns being compatible with the existence of an underlying VT causing genetic defect.

*Whole Genome Sequencing of MARTHA patients*

Genomic DNA was extracted from peripheral blood, using the BioRobot EZ1 workstation. The DNA concentration was determined using the Qubit assay kit (Thermofisher).

Whole genome sequencing was performed at the Centre National de Recherche en Génomique Humaine (CNRGH, Institut de Biologie François Jacob, Evry, FRANCE). After a complete quality control, 1µg of genomic DNA was used for each sample to prepare a library for whole genome sequencing, using the Illumina TruSeq DNA PCR-Free Library Preparation Kit, according to the manufacturer's instructions. After normalisation and quality control, qualified libraries were sequenced on a HiSeqX5 instrument from Illumina (Illumina Inc., CA, USA) using a paired-end 150 bp reads strategy. One lane of HiSeqX5 flow cell was used per sample specific library in order to reach an average sequencing depth of 30x for each sequenced individual. Sequence quality parameters have been assessed throughout the sequencing run and standard bioinformatics analysis of sequencing data was based on the Illumina pipeline to generate FASTQ file for each sample. FastQ sequences were aligned on human genome hg37 using the BWA-mem program (Li and Durbin, 2009). Variant calling was performed using the GATK HaplotypeCaller (GenomeAnalysisTK-v3.3-0, <https://software.broadinstitute.org/gatk/documentation/article.php?id=4148>). Single-sample gVCFs files were then aggregated using GATK CombineGVCFs and joint genotyping calling performed by GATK GenotypeGVCFs. Recalibration was then conducted on the whole gVCF following GATK guidelines. Following GATK VQSR, we retained single nucleotide variants in the 99.5% tranche sensitivity threshold and indels in the 99% tranche sensitivity threshold for further analysis and annotated them using Annovar (Wang *et al.*, 2010) and MORFEE.

*Variant interpretation by functional studies*

*Ex vivo* functional assays were performed as described in Labrouche-Colomer et al (Labrouche-Colomer *et al.*, submitted). Briefly, we assessed the effect of *PROC* rs116169054 (c.-33C>T; NM_000312.4) and *PLAT* rs763007802 (c.-15C>T; NM_000930.5) on the protein steady-state level in HeLa cells.

To generate the wild-type constructs (WT), the human PROC and PLAT cDNA (*NM_000312.4 c.-87_c.1383 and NM_000930.5 c.-98_1686, respectively*) were cloned into pcDNA3.1/myc-His(-) plasmid (Invitrogen) between EcoRI and HindIII or BamHI and HindIII cloning sites, respectively. PCR amplification was performed by using Phusion High Fidelity (HF) DNA Polymerase (ThermoFisher) and specific primer pairs as indicated in Supp. Table 3 (below). The amplified cDNAs were cloned in-frame with a myc-his tag. Then, the PROC *c.-33C>T* and PLAT *c.-15C>T* variants were introduced into the corresponding WT plasmids by rapid-site-directed-mutagenesis by using specific primers (Supp. Table 3), the WT plasmids as templates and Phusion HF DNA Polymerase (ThermoFisher), according to the manufacturer's recommendations. The sequence of the WT and Variant (Var) plasmids were verified by Sanger sequencing (Genewiz).

To evaluate the effect of our variants at the protein level, we transfected 1,5 µg of each of the WT and Var constructs into 6-well plates of HeLa cells, in parallel with empty pcDNA3.1/myc-His(-) vector as a negative control. Each construct was transfected in duplicate in the same experiment. Transfections were performed with jetPRIME® reagent (Polyplus Transfection) according to the manufacturer's recommendations. Forty-eight hours after transfection, we harvested the cells and extracted the total protein from each well by using the RIPA buffer (140 µl/well).

Protein concentration was determined by the BCA method (Pierce™ BCA Protein Assay Kit) and Equal amounts (50 µg) of the extracted proteins were analyzed by SDS-PAGE on 10% gels. Proteins were then transferred onto PVDF membranes (Immobilon-P or Immobilon PSQ, Merck Millipore) for immunostaining. To visualize our tagged proteins, we probed the membranes with either monoclonal anti-(c-Myc) antibody (Merck Millipore) or anti-β-actin antibody (Cell signaling technology), used here as a control. After incubation with goat anti-mouse IgG Alexa Fluor 700 (ThermoFisher), or Goat anti-Rabbit IgG (H+L) Alexa Fluor 750 (Invitrogen) secondary antibodies, blots were scanned on Odyssey Infrared Imaging System (Li-Cor Biosciences) in the 700 or 800 channels, respectively.

Germain,M. *et al.* (2011) Genetics of venous thrombosis: insights from a new genome wide association study. *PLoS ONE*, **6**, e25581.

Labrouche-Colomer,S. *et al.* (submitted) A novel rare c. -39C>T mutation in the PROS1 5’UTR causing PS deficiency by creating a new upstream translation initiation codon and inhibiting the production of the natural protein. *BIORXIV/2020/007328*.

Li,H. and Durbin,R. (2009) Fast and accurate short read alignment with Burrows-Wheeler transform. *Bioinformatics*, **25**, 1754–1760.

Oudot-Mellakh,T. *et al.* (2012) Genome wide association study for plasma levels of natural anticoagulant inhibitors and protein C anticoagulant pathway: the MARTHA project. *Br. J. Haematol.*, **157**, 230–239.

Wang,K. *et al.* (2010) ANNOVAR: functional annotation of genetic variants from high-throughput sequencing data. *Nucleic Acids Res.*, **38**, e164.

**Supplemental Table 3.** Cloning and site directed mutagenesis primers used in this study. Cloning sites are underlined. 5’P, 5’ phosphorylated primers.

| Cloning | PROC-EcoRI-F | tgcagaattcAACTCGAACTCCAGGCTGTC |
| --- | --- | --- |
|  | PROC-HindIII-R | gcccaagcttAGGTGCCCAGCTCTTCTG |
|  | PLAT-BamHI-F | tagtggatccAGAGCTGAGATCCTACAGGAG |
|  | PLAT-HindIII-R | gcccaagcttCGGTCGCATGTTGTCACG |
| Site-directed mutagenesis | PLAT-c-15CT-pF | 5’P-ATTTAAGGGATGCTGTGAAGC |
|  | PLAT-mutdir-pR | 5’P-TCACGGCTTGCTCCTTCCC |
|  | PROC-c-33CT-pF | 5’P-GTATCTCCAtGACCCGCCCCT |
|  | PROC-mutdir-pR | 5’P-TGCAAGTTCGCCGTCCTGC |

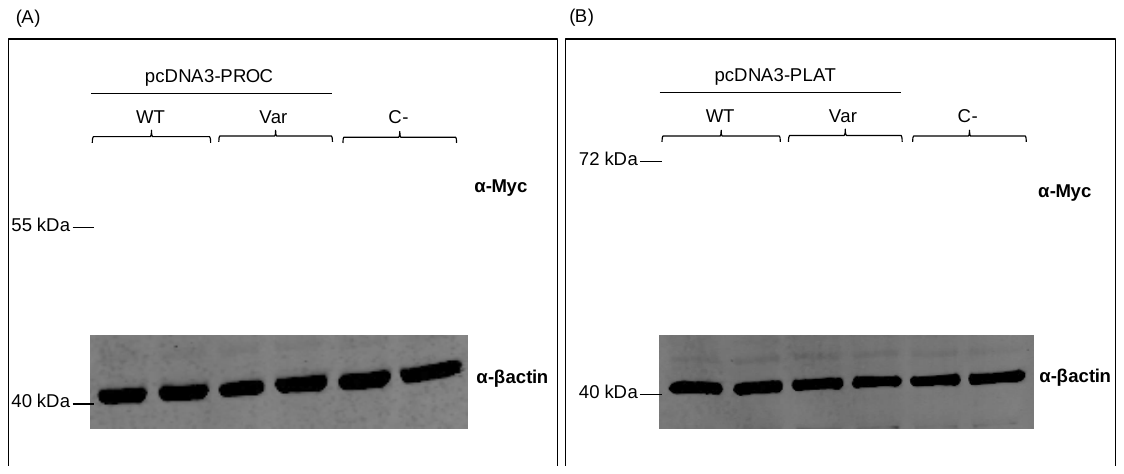

**Supplemental Figure 1.** **Evaluation of the effect of *PROC* and *PLAT* variants on the protein level.** Plasmids containing the 5’UTR and the CDS of PROC (A) or PLAT (B), in frame with a Myc-His tag, were transfected into HeLa cells in the wild-type (WT) or variant (Var) context, in parallel with an empty vector used as a negative control (C-). Whole cell proteins were extracted 48 hours after transfection and equal amounts were analyzed by western blot on 10% gels to compare the protein steady-state levels in the wild-type and variant contexts. After the transfer onto PVDF membranes, we targeted our tag-fused proteins with the monoclonal anti-(c-Myc) antibody (α-Myc) and used the anti-β-actin antibody (α-βactin) as control. Then, we scanned the membranes on Odyssey Infrared Imaging System (Li-Cor Biosciences) after incubation with the corresponding fluorescent secondary antibodies. The protein weights are indicated. kDa, kilodaton.
